# TRIB3 silencing promotes the downregulation of Akt pathway and PAX3-FOXO1 in high-risk rhabdomyosarcoma

**DOI:** 10.1101/2023.12.01.569530

**Authors:** Gabriel Gallo-Oller, Guillem Pons, Julia Sansa-Girona, Natalia Navarro, Patricia Zarzosa, Lia García-Gilabert, Paula Cabré Fernandez, Gabriela Guillén Burrieza, Lorena Valero-Arrese, Miguel F. Segura, José M. Lizcano, José Sánchez de Toledo, Lucas Moreno, Soledad Gallego, Josep Roma

**Author notes:** Correspondence: Josep Roma.

## Abstract

**Background:** Rhabdomyosarcoma (RMS), such as other childhood tumors, has witnessed treatment advancements in recent years. However, high-risk patients continue to face poor survival rates, often attributed to the presence of the PAX3/7-FOXO1 fusion proteins, which has been associated with metastasis and treatment resistance. Despite efforts to directly target these chimeric proteins, clinical success remains elusive. In this study, the main aim was to address this challenge by investigating regulators of FOXO1. Specifically, we focused on TRIB3, a potential regulator of the fusion protein in RMS.

**Methods:** TRIB3 expression was examined through the analysis of patient datasets, including gene expression profiling and gene set enrichment analyses. In cell lines, the DepMap dataset for RMS was utilized alongside Western blot analysis to assess TRIB3 expression. The functional significance of TRIB3 in RMS was assessed through constitutive and inducible shRNA-mediated knockdowns. Subsequent *in vitro* and *in vivo* analyses, including orthotopic tumor models in immune-compromised mice, were conducted to delineate the role and underlying molecular mechanisms exerted by TRIB3 in RMS

**Results:** Our findings revealed a prominent TRIB3 expression in RMS tumors, highlighting its correlation with several clinical features. By conducting TRIB3 genetic inhibition experiments, we observed an impairment on cell proliferation. Notably, the knockdown of TRIB3 led to a decrease in PAX3-FOXO1 and its target genes at protein level, accompanied by a reduction in the activity of the Akt signaling pathway. Furthermore, TRIB3 influenced posttranslational modifications, such as phosphorylation together with proteasomal degradation of PAX3-FOXO1 protein. Additionally, inducible silencing of TRIB3 significantly delayed tumor growth and improved overall survival *in vivo*.

**Conclusions:** Based on our comprehensive analysis, we propose that TRIB3 holds therapeutic potential for treating the most aggressive subtype of RMS. The findings herein reported contribute to our understanding of the underlying molecular mechanisms driving RMS progression and provide novel insights into the potential use of TRIB3 as a therapeutic intervention for high-risk RMS patients.

## Introduction

Rhabdomyosarcoma (RMS) stands as the most prevalent soft tissue malignant tumor in childhood, encompassing 4-5% of all childhood cancers [1,2]. Despite its categorization as a rare disease, RMS constitutes nearly half of all diagnosed cases of pediatric soft-tissue sarcomas [1,3]. Over the past few decades, treatment advancements have improved outcomes for RMS patients. However, patients classified as high-risk still exhibit a survival rate of only 40% [4,5]. The histologic classification of RMS has traditionally played a pivotal role in risk stratification, with alveolar RMS (ARMS) being the subtype associated with an unfavorable prognosis, followed by embryonal RMS (ERMS), which is the most common subtype [3,6]. However, in recent years, there has been a notable shift in focus towards molecular markers in the risk stratification of RMS. Specifically, two distinct genotypes have been identified: 1) those with *PAX3-FOXO1* or *PAX7-FOXO1* gene fusions (fusion positive: FP-RMS); and 2) those without these fusions (fusion negative: FN-RMS). The presence or absence of these gene fusions is currently considered a critical prognostic factor in RMS, with FP-RMS tumors exhibiting the highest rates of metastatic progression and therapy failure [7,8]. In addition, the PAX3/7-FOXO1 fusion proteins play a pivotal role in RMS biology by contributing to tumor initiation, progression, and aggressiveness. These fusion proteins function as aberrant transcription factors, driving key oncogenic processes [9,10].

Given their crucial role in the etiology of the disease and their specificity, fusion proteins are regarded as promising molecular targets for RMS. Consequently, numerous studies have focused on elucidating their molecular regulation and developing inhibitory approaches for therapeutic purposes [7,11]. Both PAX3-FOXO1 and PAX7-FOXO1 proteins present challenges as molecular targets, mainly due to their subcellular localization within the cell nucleus and their transcription factor nature [9]. However, alternative strategies, such as modulating their posttranslational modifications, could be the basis for novel therapeutic interventions for RMS. Indeed, a previous study has demonstrated that phosphorylation can inactivate PAX3/7-FOXO1, affecting its protein stabilization [9]. Despite the description of several inhibition strategies, none of them has reached clinical phases. Hence, the identification of novel regulatory elements for PAX3/7-FOXO1 holds the potential to enhance treatment approaches.

Interestingly, phosphorylation-mediated activation or inhibition, in coordination with ubiquitination, are well-documented processes which have been involved in the regulation of FOXO1 in various tumors [12–14]. Within this context, a FOXO1 key regulator is the tribbles homolog 3 protein (TRIB3), which plays a pivotal role in diverse malignancies, including breast cancer and hepatocellular carcinoma [15,16]. TRIB3 belongs to a subfamily of pseudokinases comprising three members: TRIB1, TRIB2, and TRIB3. TRIB proteins share a highly conserved, non-functional kinase-like domain and appear to have evolved two major mechanisms of action: one related to E3 ligase-dependent ubiquitination and the second as a scaffolding function, involved in binding and regulating protein kinases [17]. Notably, the TRIB3/Akt/FOXO1 regulatory axis has been implicated in various malignancies and diseases [15,16,18]. While the role of TRIB3 has been studied in several tumor types, its involvement in RMS remains unexplored. Of note, a precedent study by our team [19] pointed TRIB3 as a possible regulator of chemoresistance in RMS, suggesting that TRIB3 may play significant functional roles in this tumor.

In this work, we present the first functional characterization of the role of TRIB3 in RMS. Our results prove that TRIB3 silencing causes a substantial impairment on RMS cell proliferation, along with modulation of the PAX3-FOXO1 activity and a downregulation of the Akt signaling pathway. Collectively, these findings not only shed light on the previously unexplored role of TRIB3 in RMS but also introduce a novel therapeutic avenue for high-risk patients. Further investigation into the potential of TRIB3 as a therapeutic target in RMS holds great promise for refining treatment approaches and ultimately improving outcomes for patients facing this challenging disease.

## Results

### TRIB3 is overexpressed in RMS and correlates with the presence of the fusion protein

To investigate the oncogenic role of the TRIB family in RMS, we conducted a transcriptomic data mining of multiple RMS datasets. The expression levels of TRIB family members were examined in relation to several clinical features, including histology, disease stage, and the presence or absence of fusion proteins.

Our analysis revealed varied expression patterns among the TRIB family members in RMS. Notably, *TRIB3* displayed high expression levels in RMS samples relative to muscle tissue. This increased expression was also evident in the ARMS subtype in contrast to the ERMS subtype, and also at stage 4 of the disease (Figure 1A-C). Moreover, FP-RMS tumors (both PAX3-FOXO1 and PAX7-FOXO1) exhibited elevated *TRIB3* expression compared to FN-RMS (Figure 1D). In contrast, *TRIB1* showed reduced expression in RMS tumors compared to healthy muscle tissue, and this was consistently observed in both the ARMS subtype and FP-RMS tumors (Supplementary Figure 1A). Interestingly, while *TRIB2* expression was higher compared to the healthy counterpart, the ARMS subtype exhibited a low *TRIB2* expression relative to the ERMS subtype. This lower *TRIB2* expression was also observed at the advanced disease stage (stage 4), as well as in the context of FP-RMS *vs* FN-RMS (Supplementary Figure 1B).

**Figure 1.**
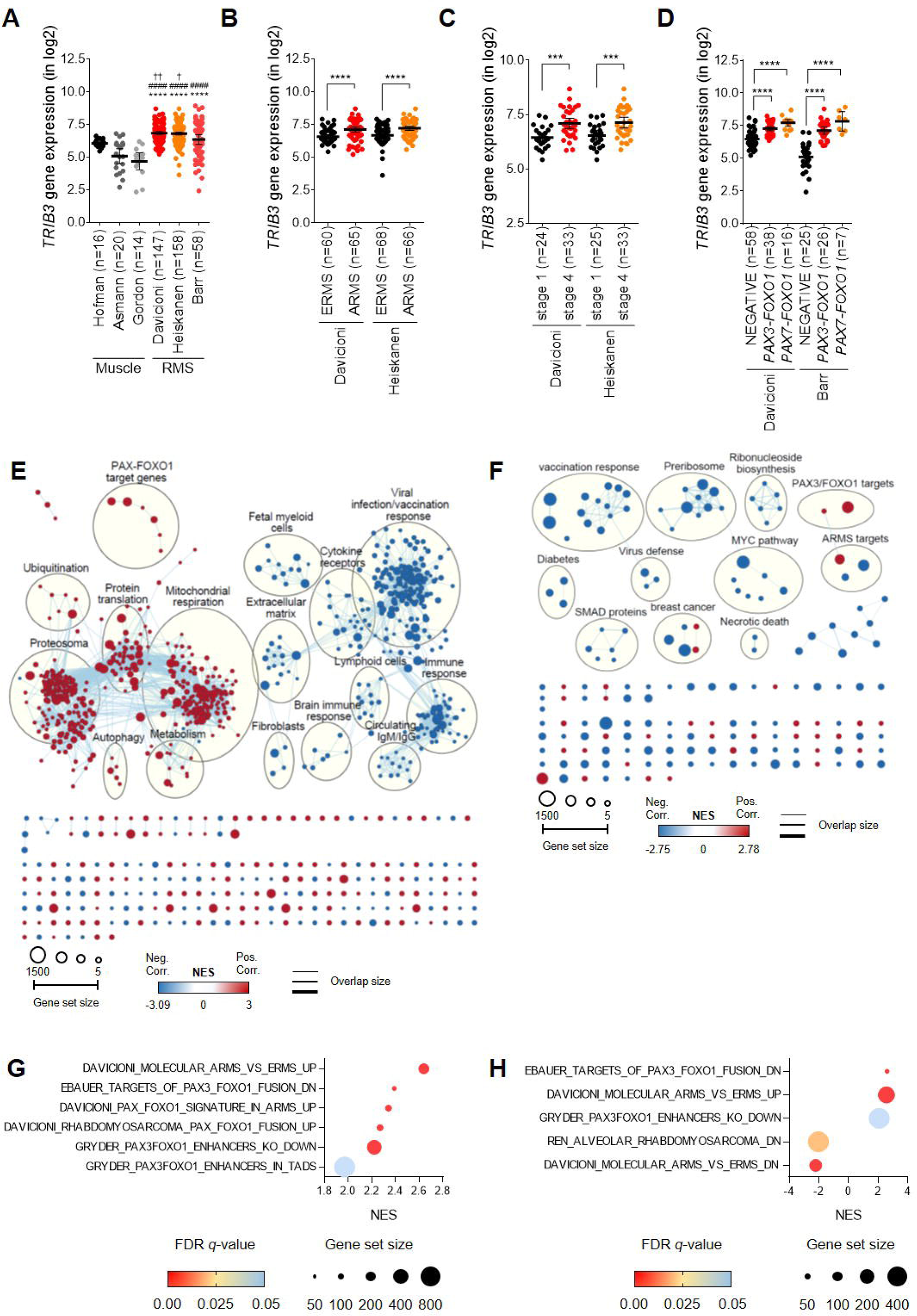
Analysis of TRIB3’s role and its regulatory network in RMS. Expression data obtained from patients’ datasets on the R2 platform were correlated with several clinical features. The data are presented as the mean with 95% confidence interval (CI). (A) TRIB3 was found to be overexpressed in rhabdomyosarcoma datasets compared to healthy counterparts. One-way ANOVA was performed, and Tukey’s multiple comparisons test was used as the post hoc test. Significance versus Hofman is indicated by †, versus Asmann by #, and versus Gordon by *. (B) The ARMS subtype exhibited the highest TRIB3 expression levels. Unpaired t-tests were conducted for each dataset. (C) TRIB3 showed significant overexpression in late disease stages (unpaired t-test). (D) Tumors harboring PAX3-FOXO1 or PAX7-FOXO1 fusion gene displayed higher TRIB3 expression levels. One-way ANOVA with Dunnett’s post hoc test was used to determine significance. (E) Gene Set Enrichment Analysis (GSEA) using the Barr patients’ dataset revealed an enrichment map visualized by Cytoscape, demonstrating gene sets significantly enriched in FP-RMS patients (n=25). (F) Similarly, gene sets significantly enriched in RMS cell lines were also visualized through the enrichment map. In the network map, node size represents the number of genes in the gene-set; edge thickness is proportional to the overlap between gene-sets and the enrichment score is mapped to the node color as a color gradient. (G) Bubble plots graphically represent gene sets specifically associated with the fusion protein and significantly enriched in FP-RMS patients, while (H) they similarly depict gene sets enriched in RMS cell lines. FDR q-value is represented by the color of the circles while the size of the circles represents the number of identified genes within each gene set. NES: normalized enrichment score.

### Regulatory network analysis identifies TRIB3 as a putative PAX3-FOXO1 regulator

Considering the lack of studies of TRIB3 in RMS, and to elucidate the possible molecular mechanism, we conducted a pre-ranked Gene Set Enrichment Analysis (GSEA) using gene expression data from patient samples and RMS cell line datasets together with Cytoscape visualization software. GSEA in FN-RMS patient samples revealed enrichment in gene sets associated with the tricarboxylic acid cycle (TCA) and unfolded protein response (UPR) (Supplementary Figure 1C). In the FP-RMS group, enrichment in gene sets related to previously described TRIB3-associated mechanisms, including metabolism, cell energy, ubiquitination, immune system, and viral infection were detected (Figure 1E). In addition, GSEA using expression data from 11 RMS cell lines revealed similar results as observed in patient datasets (Figure 1F).

However, the most significant and remarkable enrichments were observed in gene sets directly related to the ARMS expression signature and PAX3/7-FOXO1 target genes, both in patients’ datasets (Figure 1G and Supplementary Figure 1D) and cell lines (Figure 1H and Supplementary Figure 1E). Notably, among the enriched gene sets was DAVICIONI-MOLECULAR-ARMS-VS-ERMS-UP, which contains genes associated with the molecular distinction between ARMS and ERMS subtypes. Other significantly enriched gene sets were directly linked to the downstream effects of PAX3-FOXO1 and its target genes, such as GRYDER-PAX3FOXO1-ENHANCERS-KO-DOWN and GRYDER-PAX3FOXO1-ENHANCERS-IN-TADS, which include genes related to molecular processes regulated by PAX3-FOXO1. Additionally, the EBAUER-TARGETS-OF-PAX3-FOXO1-FUSION-DN and DAVICIONI-RHABDOMYOSARCOMA-PAX-FOXO1-FUSION-UP gene sets were significantly enriched, consisting of gene targets regulated by PAX3-FOXO1.

Our analysis demonstrates enrichments in gene sets closely linked to the ARMS expression signature, as well as the PAX3/7-FOXO1 and its downstream effects and target genes in FP-RMS. This observation strongly suggests a significant role for TRIB3 within the critical molecular pathways involved in RMS oncogenesis.

### TRIB3 is overexpressed in FP-RMS cell lines

Next, TRIB3 protein expression was characterized in a panel of RMS cell lines (Figure 2A). Consistent with the findings in patient samples, the ARMS cell lines exhibited the higher levels of TRIB3 protein expression than the ERMS cells, except for the CW9019 cell line (positive for PAX7-FOXO1) (Figure 2B). Moreover, TRIB3 protein expression was significantly higher in PAX3-FOXO1 positive cells (RH4, RH30 and RH28) than in FN-RMS cell lines (RH18, RD, RH36 and RUCH-2) (Figure 2C). Taken together, these results suggest a potential functional role of TRIB3 in FP-RMS tumors, particularly those carrying the PAX3-FOXO1 variant.

**Figure 2.**
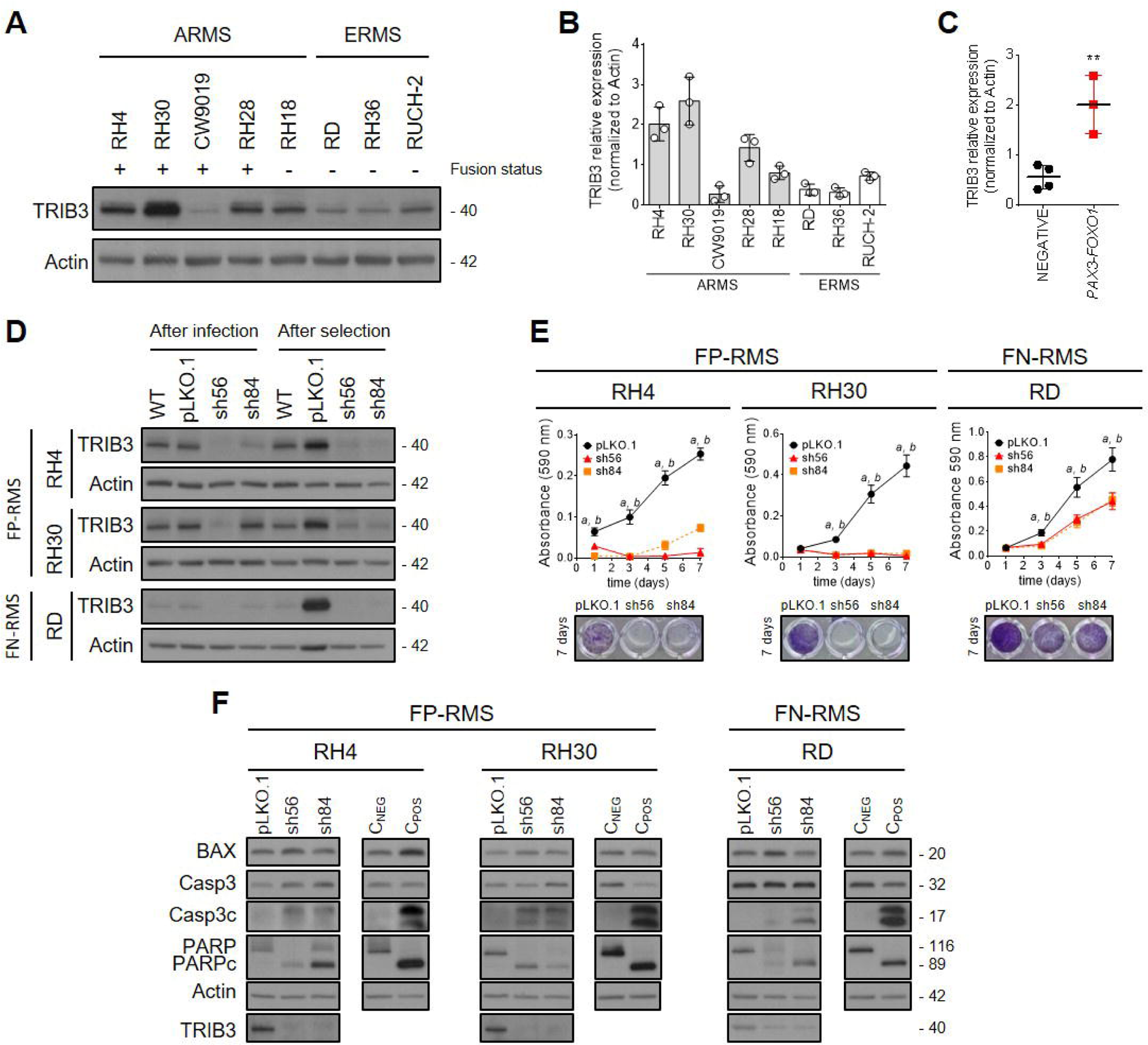
TRIB3 is overexpressed in FP-RMS cell lines, and its genetic inhibition leads to impaired cell survival and increased apoptosis. (A) TRIB3 protein expression was assessed by WB in RMS cell lines. The fusion status of each cell line is indicated by + for FP-RMS cell lines and - for FN-RMS. (B) Except for the CW9019 cell line, cell lines classified as ARMS exhibited the highest TRIB3 protein levels. (C) Fusion-positive cell lines demonstrated the highest TRIB3 protein levels. (D) To investigate the potential role of TRIB3, a constitutive model for genetic inhibition was optimized. sh56 and sh84 were selected to silence TRIB3 in RH4, RH30 (both FP-RMS), and RD (FN-RMS) cell lines. TRIB3 protein levels were determined 24 h after lentiviral infection and 72h after selection with puromycin. (E) Following constitutive TRIB3 knockdown, a clear impairment in cell proliferation was observed in FP-RMS cell lines compared to FN-RMS cell lines. (F) Analysis of apoptotic markers by WB confirmed the induction of apoptosis after TRIB3 silencing.

### TRIB3 silencing impairs proliferation and induces apoptosis in RMS cell lines

To characterize the functional role of TRIB3 in RMS models, we conducted loss of function experiments using shRNA-mediated silencing of TRIB3. We tested five different shRNA constructs targeting TRIB3 in the RH30 cell line, the model that exhibited the highest TRIB3 protein levels (Supplementary Figure 2A). Two shRNA, namely sh56 and sh84, were selected for further functional experiments. Considering the possible association between TRIB3 and PAX3-FOXO1, we chose the cell lines RH4 and RH30, which displayed high TRIB3 levels and are derived from FP-RMS. Additionally, we included the RD cell line as a control (derived from FN-RMS and having low TRIB3 protein levels). TRIB3 knockdown was performed in all three cell lines, and a significant reduction in TRIB3 expression was confirmed after selection with puromycin (Figure 2D and Supplementary Figure 2B for blot quantification).

Proliferation assays revealed that, upon TRIB3 knockdown, FP-RMS cells (RH4 and RH30) showed no ability to proliferate (Figure 2E). In contrast, the TRIB3 knockdown in the FN-RMS cell line RD, partially impaired proliferation. Analysis of apoptotic markers (Figure 2F) also revealed that TRIB3 knockdown induced apoptosis in the three cell lines, as shown by increased levels of cleaved caspase-3 and cleaved PARP (blot quantification in Supplementary Figure 2C). Taken together, these results provide evidence that TRIB3 silencing induces apoptotic cell death in RMS cells.

### Inducible TRIB3 genetic inhibition confirmed proliferation impairment and apoptosis induction in RMS

To validate the findings obtained with the constitutive model, sh56 and sh84 shRNAs were cloned into the pLKO-3xLacO IPTG-inducible system. The three cell lines in this study (RH4, RH30, and RD) were transduced with this system, and TRIB3 protein levels were assessed by Western blot (WB) at different concentrations of IPTG. A decrease in TRIB3 protein levels was observed in a concentration-dependent manner for the three cell lines analyzed (Figure 3A, and Supplementary Figure 3A for blot quantification). Furthermore, the reduction in cell survival exhibited a pronounced dependence on the concentration of IPTG, with this reliance being particularly evident 7 days post-induction (Figure 3B and Supplementary Figure 3B). This observation implies a significant correlation between TRIB3 levels and the observed effects. Additionally, these results align with the findings in the constitutive model, further supporting the notion that FP-RMS cells exhibit higher sensitivity to TRIB3 silencing. The determination of apoptotic markers showed similar results to those observed with the constitutive model. After 96h in the presence of 50µM IPTG, an increase in cleaved caspase 3 was observed with both shRNAs. Furthermore, the results indicated a decrease in total levels of PARP in the RH4, RH30, and RD cell lines. Specifically, cleaved PARP was only detected in the RH4 and RH30 cell lines, while the RD cell line exhibited a reduction in total PARP without any observable cleaved form (Figure 3C, with blot quantification in Supplementary Figure 3C).

**Figure 3.**
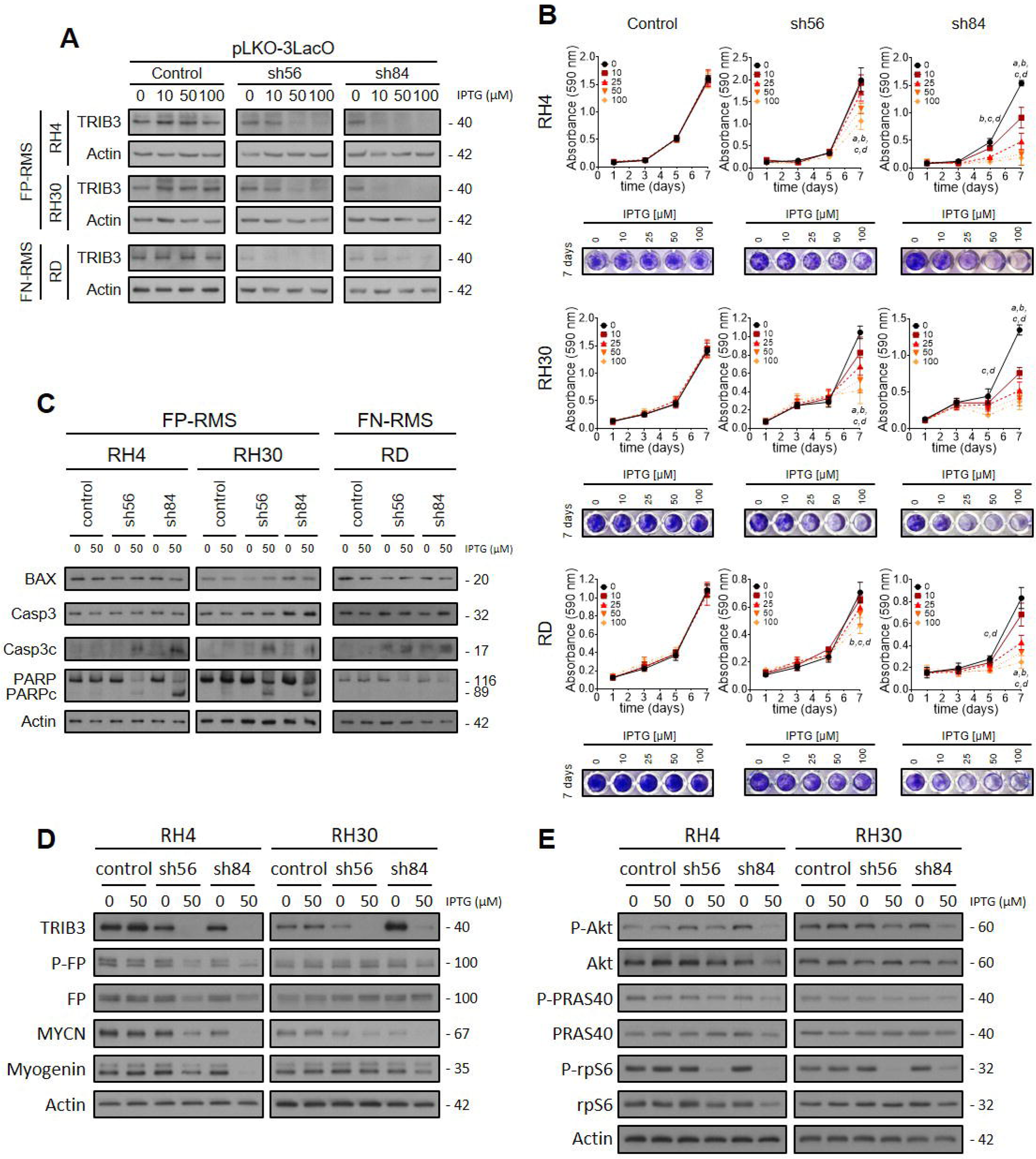
TRIB3 levels modulate proliferation by inducing apoptosis through the regulation of PAX3-FOXO1 activity, along with modulation of the Akt signaling pathway. sh56 and sh84 were cloned into the IPTG-inducible pLKO-3xLacO plasmid. (A) TRIB3 genetic inhibition in a concentration-dependent manner was confirmed in the 3 cell lines. (B) Proliferation was assessed using the crystal violet assay at different IPTG concentrations. A two-way ANOVA followed by Dunnett’s multiple comparisons test was performed to compare each IPTG concentration with the non-induced condition at each time point. (C) Apoptosis was analyzed by determining apoptotic markers through WB. (D) Analysis of fusion protein and its phosphorylation status by WB, along with MYCN and myogenin proteins as fusion protein target genes, confirmed that PAX3-FOXO1 function is affected after TRIB3 knockdown. (E) Akt signaling pathway inhibition after TRIB3 knockdown was analyzed by WB. Phosphorylation status of Akt and its main downstream effectors (PRAS40 and rpS6) was assessed.

### TRIB3 genetic inhibition reduced the expression of fusion protein targets and impaired Akt signaling pathway

To investigate the influence of TRIB3 silencing on PAX3-FOXO1 protein, we evaluated the PAX3-FOXO1 protein level and its phosphorylation (at the Ser503 site) in RH4 and RH30 cell lines after TRIB3 knockdown. In RH4 cells, we observed a reduction in the total protein levels concomitant with a noticeable decrease in Ser503 phosphorylation upon TRIB3 silencing. In contrast, total PAX3-FOXO1 remained unaffected in RH30 cells. Remarkably, the levels of fusion protein targets, MYCN and myogenin, were decreased in both cell lines following TRIB3 knockdown (Figure 3D, and Supplementary Figure 3D for blot quantification).

Previous studies have underscored the importance of TRIB3 in modulating the Akt signaling pathway and its cooperative contribution to oncogenic processes [16,20]. Hence, we sought to investigate the impact of TRIB3 on the Akt pathway in FP-RMS cell lines. Following TRIB3 knockdown, both RH4 and RH30 cell lines displayed a reduction in Akt phosphorylation (Ser473), together with a reduction in the phosphorylation of downstream effectors such as PRAS40 (at Ser235/236) and rpS6 (Ser235/236) (Figure 3E, and Supplementary Figure 3E for blot quantification). These results suggest that TRIB3 is essential for maintaining the activity of pro-survival signals such as Akt pathway.

### TRIB3 interacts with PAX3-FOXO1 and Akt in RMS

Interactions between TRIB3 and Akt have been reported in various contexts [16,21,22], suggesting a significant role of TRIB3 within the regulatory axis of TRIB3/Akt/FOXO1 [15,16,18,23]. However, although one study preliminary describes TRIB3 as a potential regulator of PAX3-FOXO1 in RMS [9], this interaction have not been explored in RMS. To clarify this, we next sought to investigate whether TRIB3 interacts with PAX3-FOXO1 and/or Akt in FP-RMS cell lines. We assessed the interaction of TRIB3 with PAX3-FOXO1 or Akt by co-immunoprecipitation of proteins derived from RH4 or RH30 cell lysates. PAX3-FOXO1 and Akt were co-immunoprecipitated with TRIB3 (Figure 4A), indicating that TRIB3 interacts with both PAX3-FOXO1 and Akt in RMS cells.

**Figure 4.**
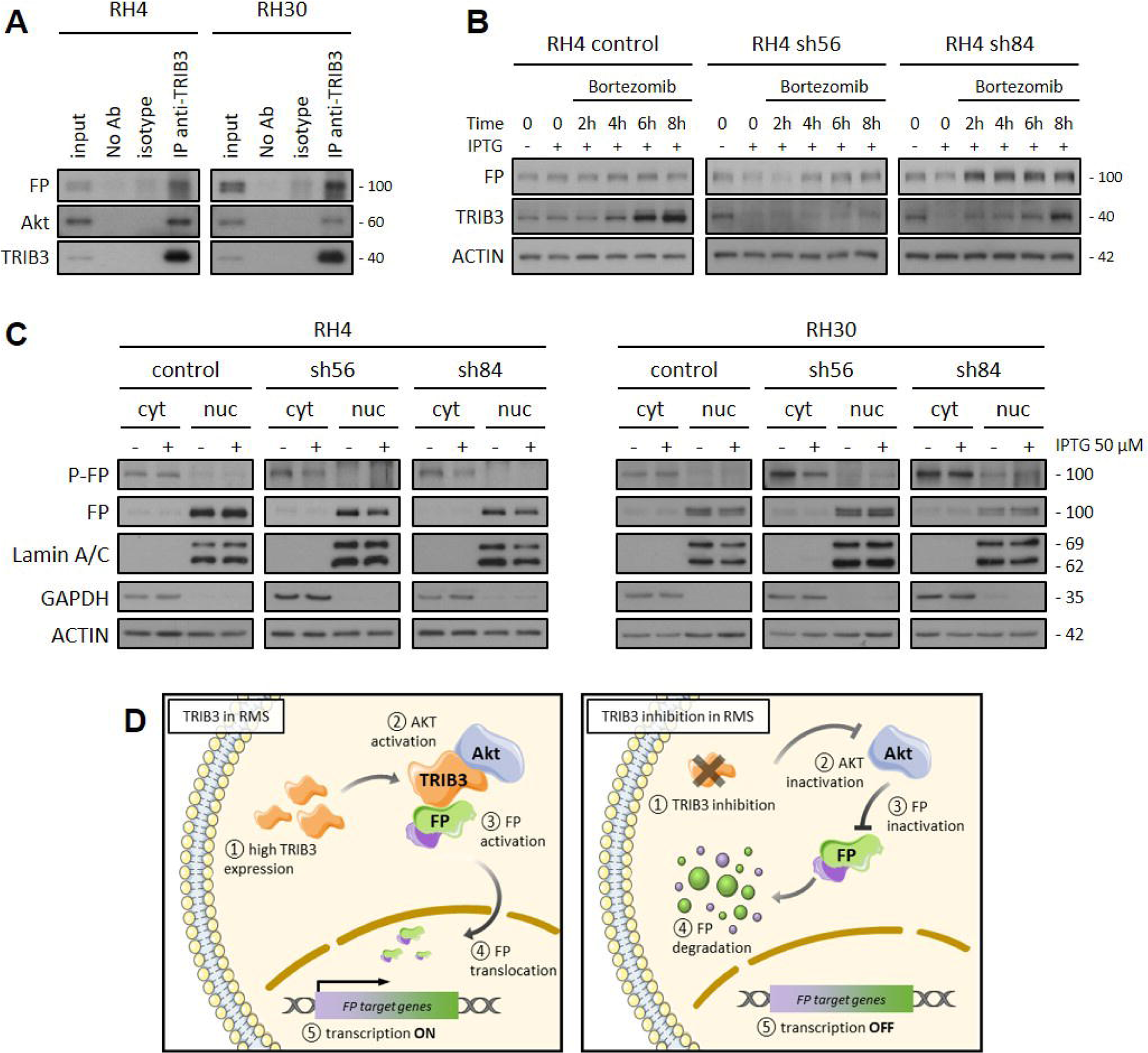
TRIB3 interacts with Akt and PAX3-FOXO1 modulating PAX3-FOXO1 stability via proteasome-mediated degradation and cytoplasmic phosphorylation in FP-RMS cells. (A) WB of co-immunoprecipitation of Akt and PAX3-FOXO1 after TRIB3 immunoprecipitation in RH4 and RH30 cells. Negative control was included (beads only and no antibody), and isotype-matched IgG was used as a control. (B) Treatment with Bortezomib led to a rescue in PAX3-FOXO1 total levels in the RH4 cell line after TRIB3 silencing. RH4 cells were transfected with either the control plasmid or specific shRNA constructs (sh56 or sh84) targeting TRIB3. The cells were induced for 96 hours by the addition of IPTG to the media, followed by bortezomib treatment at 25 nM. Subsequently, cells were harvested at various time points and subjected to Western blot analysis to determine the levels of PAX3-FOXO1 and TRIB3. (C) Cytoplasmic and nuclear fractions were analyzed by WB after TRIB3 knockdown in RH4 and RH30 cell lines. (D) Proposed molecular mechanism by which TRIB3 can modulate PAX3-FOXO1 stability is as follows: TRIB3, in conjunction with Akt, interacts with the fusion protein, thereby modulating its phosphorylation and promoting stabilization against proteasome degradation. Upon TRIB3 silencing, phosphorylation of the fusion protein is lost, leading to degradation by the proteasome.

### TRIB3 enhances the phosphorylation of and proteasome-mediated of PAX3-FOXO1

To investigate the mechanism underlying the decrease in total levels of PAX3-FOXO1 after TRIB3 silencing, we explored the effect of proteasome inhibitor bortezomib. In the RH4 cells transfected with the control plasmid, bortezomib induced a gradual increase in PAX3-FOXO1 and TRIB3 levels over time (Figure 4B, and Supplementary Figure 4A for blot quantification), suggesting that proteasome degradation plays a role in the decrease of PAX3-FOXO1. Moreover, cells harboring sh56 or sh84 and induced by IPTG were also treated with bortezomib. Interestingly, a rescue in the levels of the PAX3-FOXO1 was observed over time in cells transfected with sh56 and sh84, indicating that proteasome inhibition partially restored PAX3-FOXO1 levels.

Finally, we investigated the molecular mechanism associated with the phosphorylation of PAX3-FOXO1 by analyzing separate cytoplasmic and nuclear protein fractions in RH4 and RH30 cell lines (Figure 4C, and Supplementary Figure 4B for blot quantification). Upon TRIB3 silencing in RH4 cells, a decrease in the total levels of the fusion protein was specifically observed in the nuclear fraction of cells with sh56 and sh84. Nonetheless, in the RH30 cell line, there was no discernible alteration in the overall levels of the fusion protein in either the cytoplasmic or nuclear fractions. Interestingly, a striking reduction in the phosphorylation levels was consistently observed only in the cytoplasmic fraction for both sh56 and sh84, regardless of the cell line. These findings suggest a potential functional role of the phosphorylated form of PAX3-FOXO1 in the cytoplasm, in addition to its accepted nuclear function, challenging the conventional notion of its exclusive nuclear functionality.

### TRIB3 knockdown impairs tumor growth in vivo

Finally, we evaluated the impact of TRIB3 genetic inhibition in an *in vivo* model. RH30 cells transduced with either the pLKO-3xLacO control plasmid or the IPTG-inducible shTRIB3 construct (sh84) were injected into the gastrocnemius muscle of SCID mice. Once the primary tumors were detected, mice were randomized into vehicle (IPTG -) and IPTG receiving groups (IPTG +). The growth of control-vector transduced tumor cells was not affected by the administration of IPTG and no significant differences in mice survival were observed (Figure 5A). TRIB3 protein levels were also not modified in the control group (Figure 5B). Conversely, in shTRIB3 group, a significant delay in tumor growth and improved overall survival were observed in the presence of IPTG (Figure 5C). When primary tumors were analyzed at the end of the experiment, the TRIB3-silenced group exhibited a notable reduction of approximately 50% in TRIB3 protein levels (Figure 5D).

**Figure 5.**
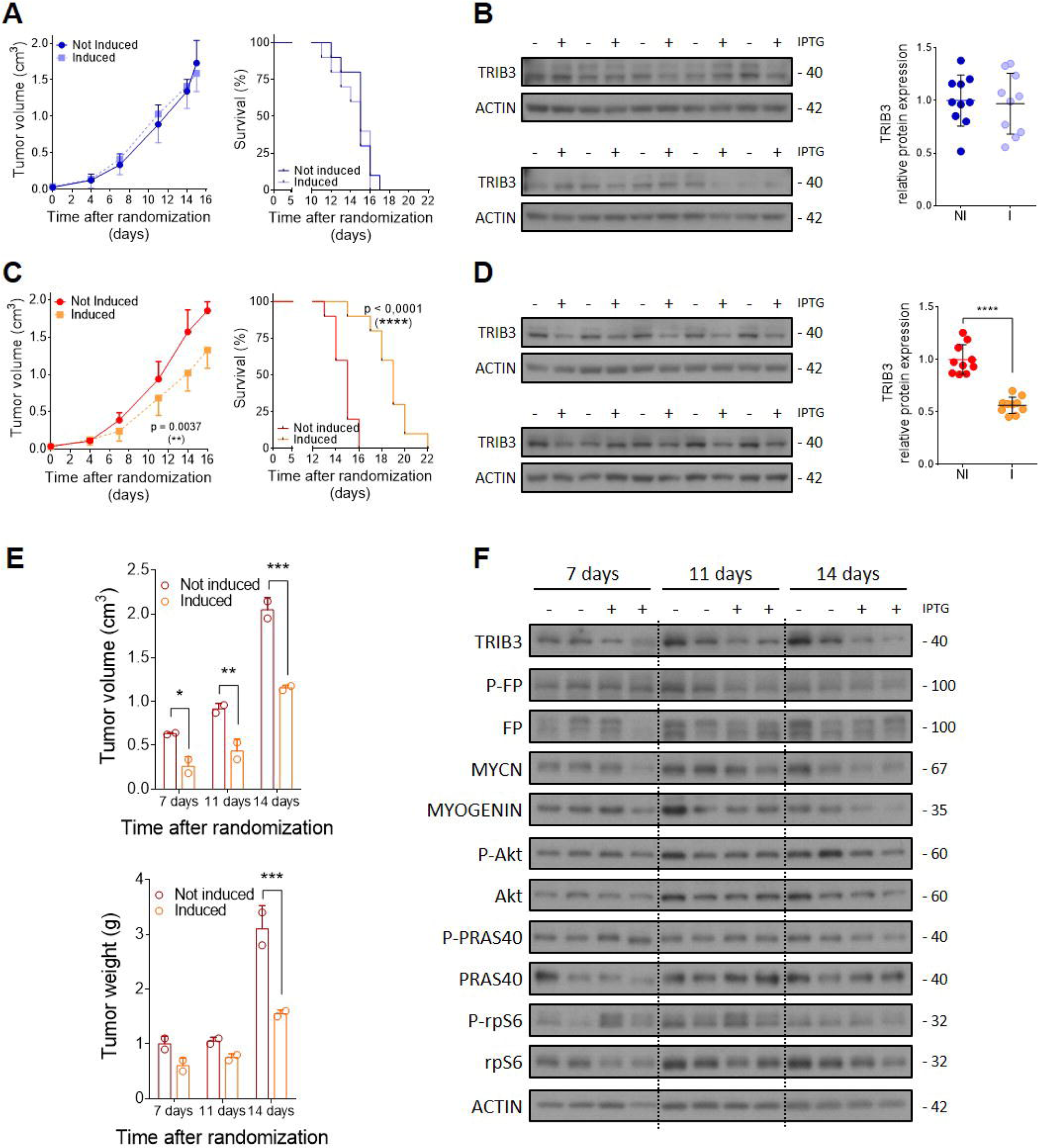
Suppression of TRIB3 inhibits *in vivo* tumor growth. An *in vivo* murine assay was conducted using the inducible pLKO-3xLacO system in the RH30 cell line. (A) Tumor growth (left panel) and survival (right panel) of control plasmid-derived tumors showed no significant differences after induction. (B) TRIB3 protein levels in tumors were analyzed by WB (left panel) and quantified (right panel). (C) Induction of sh84 in tumor cells resulted in delayed tumor growth (left panel) and improved survival (right panel). (D) WB analysis revealed decreased TRIB3 protein levels after IPTG induction (left panel), with a significant reduction of approximately 50% observed in induced tumors (right panel). (E) Mice from the sh84 group, sacrificed at 7, 11, and 14 days, showed a significant decrease in both tumor volume (upper panel) and tumor weight (lower panel). (F) Analysis of tumor samples by WB showed a decrease in MYCN and myogenin levels, along with decreases in total Akt levels and in the phosphorylation status of its substrate PRAS40. In (B) and (D) each lane represents a sample from an individual mouse.

In order to validate the effect of TRIB3 silencing on downstream effectors identified *in vitro*, mice were sacrificed at different time points during the experiment (Supplementary Figure 5A). At 7-, 11- and 14-days post-induction of TRIB3 silencing, a decrease in tumor volume and tumor weight was observed (Figure 5E). WB analysis confirmed TRIB3 inhibition. Regarding the downstream effectors, no decrease in phosphorylation or fusion protein total levels were observed. On the contrary, a decrease in PAX3-FOXO1 target genes MYCN and myogenin was observed after 14 days of induction with IPTG. Regarding Akt signaling pathway, only a decrease in total Akt levels was observed at 14 days, along with a decrease in PRAS40 phosphorylation (Ser235/236) at the same time point (Figure 5F and Supplementary Figure 5B for blot quantification).

## Discussion

Rhabdomyosarcoma (RMS) remains a therapeutic challenge, particularly for high-risk patients [24,25]. Targeting PAX3/7-FOXO1, a hallmark of RMS, has proven to be exceedingly difficult, adding complexity to the search for effective therapies [26,27]. Understanding the molecular mechanisms underlying RMS pathogenesis and the regulation of fusion protein is crucial for developing targeted treatments. In this study, we investigated the role of TRIB3 in RMS and explored its potential functional associations and molecular mechanisms related to PAX3-FOXO1.

One of our key findings was the significant overexpression of TRIB3 in RMS, particularly in RMS patients and cells carrying PAX3-FOXO1 or PAX7-FOXO1 fusion proteins. These findings align with previous research that showed elevated TRIB3 expression in several cancers [28–30], suggesting that TRIB3 might have pro-oncogenic functions in RMS. This potential role could contribute to a poorer prognosis and increased resistance to chemotherapy and radiotherapy [29,31–33]. Additionally, we identified a significant interaction between TRIB3 and fusion protein, implying that TRIB3 likely participates in the regulation of PAX3-FOXO1 in RMS. While this association had been previously proposed in RMS through *in silico* analysis [9], experimental confirmation was lacking. However, a similar interaction between TRIB3 and FOXO1 has been reported in breast cancer [16], highlighting the significance of investigating factors that could modulate FOXO1 as valuable tools in identifying potential regulators of PAX3/7-FOXO1 in RMS.

To validate the functional significance of TRIB3, we conducted genetic inhibition experiments in RMS cell lines using both constitutive and inducible systems. The consistency of the results between these two distinct experimental approaches provides robust evidence for the critical role of TRIB3 in RMS. Silencing of TRIB3 led to impaired proliferation and increased apoptosis in fusion-positive RMS cells, highlighting their dependency on TRIB3. These observations align with prior studies that have demonstrated that TRIB3 silencing induces apoptosis and reduces cell proliferation in various types of cancer cells [32,34–36]. Conversely, there are reports suggesting the opposite effect, where overexpression of TRIB3 yields opposite results in other cancer types [15,37]. Nevertheless, our findings underscore the dual role of TRIB3 in cancer, an aspect that has been extensively reviewed [17,38,39]. Therefore, TRIB3 emerges as a promising therapeutic target for RMS, offering potential avenues for targeted interventions.

At the molecular level, we obtained intriguing results when investigating the PAX3-FOXO1 protein levels following TRIB3 knockdown. We observed a reduction in total PAX3-FOXO1 levels, implying that TRIB3 may regulate the stability and activity of the fusion protein within RMS cells. This regulation potentially involves the phosphorylation of PAX3-FOXO1, particularly at Ser503, which has been described as stabilizing of the fusion protein [9]. In fact, phosphorylation of the fusion protein is a well-documented mechanism that modulates its transcriptional activity and stability [9,40,41]. Additionally, the reduction in PAX3-FOXO1 targets, MYCN and myogenin, further supports TRIB3 potential regulatory role in downstream signaling events associated with tumor progression. In addition, TRIB3 knockdown led to impaired Akt signaling, consistent with TRIB3 known role in regulating Akt [16,42,43], a critical pathway for cancer cell survival and proliferation. The crosstalk between TRIB3 and Akt in promoting oncogenesis underscores the importance of analyzing the Akt pathway following TRIB3 knockdown. Interestingly, a regulatory axis TRIB3/Akt/FOXO1 has been described in other malignancies and diseases [15,16,18,20,23], suggesting that TRIB3’s, in cooperation with Akt, may exert an influence on FOXO1, which could also be operative in the context of PAX3-FOXO1 within RMS.

Remarkably, we found an interaction between TRIB3 and PAX3-FOXO1, as well as with Akt, indicating a potential mechanism for TRIB3 influence on the fusion protein’s activity. Additionally, TRIB3 appears to have a significant role in fusion protein stability by modulating its phosphorylation status, underscoring its importance in regulating PAX3-FOXO1 function. Indeed, regulation by the proteasomal system has been suggested previously for PAX3-FOXO1 in RMS [9,44,45], but the process and possible regulators of this process in RMS remain unclear.

Our results also yield valuable insights into the subcellular localization and phosphorylation dynamics of the fusion protein following TRIB3 silencing, suggesting TRIB3’s involvement in controlling the phosphorylation and nuclear accumulation of PAX3-FOXO1 in RMS. Furthermore, our data imply the involvement of the Akt signaling pathway in conjunction with TRIB3 as potential modulators of PAX3-FOXO1 activity. This regulatory axis, TRIB3/Akt/FOXO1, has been previously described in other malignancies, further supporting TRIB3’s potential role in RMS. However, the mechanism of action appears to be context specific. Some reports indicate that TRIB3 can activate Akt, which subsequently phosphorylates FOXO1 and other FOXO members to prevent their nuclear translocation [15,18,46,47]. In contrast, other studies suggest that the interaction between TRIB3 and Akt can activate FOXO1, allowing its nuclear accumulation [16,20]. In the case of RMS, we can hypothesize that the interaction between TRIB3 and Akt regulates PAX3-FOXO1 phosphorylation at Ser503, described as a stabilizer of PAX3-FOXO1 [9], and could also further affect its nuclear export (Figure 4D). In fact, this site is equivalent to S322 in FOXO1, which has been described as a regulator of its nuclear localization [48,49]. In summary, the apparent cytoplasmic occurrence of PAX3-FOXO1 phosphorylation in RMS cells and the interaction between TRIB3 and Akt open new avenues for targeting the fusion protein. In fact, the therapeutic potential of disrupting the interaction between TRIB3 and Akt has been tested at the preclinical level for breast and lung cancer [16,20].

Our *in vivo* experiments further confirmed the impact of TRIB3 on RMS cell proliferation and revealed its role in modulating PAX3-FOXO1 target genes and Akt signaling *in vivo*. These findings align with studies in other tumor types, such as lung adenocarcinoma [35], glioblastoma [32] and ovarian cancer [34], where TRIB3 down-regulation also impaired tumor growth. Interestingly, *in vivo* growth inhibition did not reach levels to those observed *in vitro*. A plausible explanation could be the different levels of TRIB3 knockdown observed. While very high knockdown levels were achieved *in vitro*, only a reduction of approximately 50% was observed *in vivo*. In fact, the inducible model demonstrates that growth inhibition in RMS cells occurs in a dose-dependent manner. Moreover, our observations of TRIB3’s effects on PAX3-FOXO1 target genes and the Akt signaling pathway provide valuable insights into the molecular mechanisms driving RMS development and progression in an *in vivo* setting. These results confirmed TRIB3 impact on RMS cell proliferation and its role in modulating PAX3-FOXO1 and Akt signaling, reinforcing the translational potential of TRIB3 as a therapeutic target for RMS. The development of therapeutic approaches based on the disruption of TRIB3 interactions could potentially facilitate the development of effective therapeutic strategies for ultimately addressing the ‘undruggable’ PAX3-FOXO1 in RMS.

## Conclusion

In conclusion, this study provides valuable insights into the crucial role of TRIB3 in RMS pathogenesis, particularly in PAX3-FOXO1 tumors, the most aggressive RMS subtype. The overexpression of TRIB3 in RMS, its functional impact on proliferation, apoptosis, PAX3-FOXO1 regulation, and Akt signaling support its potential as an oncogenic player and therapeutic target. These findings offer new avenues for therapeutic development and underscore the need for further research to validate TRIB3 as a promising candidate for targeted therapy in RMS. The knowledge gained from this study contributes to our understanding of the complex molecular landscape of RMS and facilitates the way for more effective treatments to improve patient outcomes.

## Material and methods

### Mining transcriptome data for rhabdomyosarcoma

We conducted an mRNA expression analysis using the R2 Genomics Analysis and Visualization platform (http://r2.amc.nl). The expression values for *TRIB1*, *TRIB2*, and *TRIB3* genes were compared based on histological, molecular, and clinical features. The following datasets were included in our analysis: 1) Normal Muscle - Hofman - 121 - MAS5.0 - u133a: only the control group was selected (n=16) [50]; 2) Normal Muscle Skeletal - Asmann - 40 - MAS5.0 - u133p2: the analysis considered only young individuals to avoid age-related bias (n=20) [51]; 3) Normal Muscle - Gordon - 22 - MAS5.0 - u133p2: only the subset of young patients (n=14) was included (n=14) [52]; 4) Tumor Rhabdomyosarcoma - Davicioni - 147 - MAS5.0 - u133a: this dataset comprised 147 tumor samples (n=147) [53]; 5) Tumor Alveolar Rhabdomyosarcoma - Heiskanen - 186 - MAS5.0 - u133a: only patient tumor samples were used for the analysis, excluding cell line data (n=158) [53]; and 6) Tumor Rhabdomyosarcoma - Barr - 58 - MAS5.0 - u133p2 (n=58) [54]. The selection of RMS datasets was based on the number of patients and the availability to subclassify patients according to different features. Among the selected datasets, the Davicioni and Heiskanen datasets provided valuable patient subclassification based on histological subtype or disease stage. Additionally, the Davicioni and Barr datasets were used to investigate the expression patterns in FP-RMS or FN-RMS tumors.

For RMS cell lines, expression values were obtained from the DepMap portal (www. https://depmap.org/portal/). The values were derived from the Expression 22Q2 Public dataset and expressed as Log2-transformed. 11 RMS cell lines were selected for the analysis, including JR, RH28, RH30, RH4, RHJT, SCMCRM2 and CW9019 (classified as ARMS); RD, SMSCTR, RMSYM and TTC442 (classified ERMS).

### Gene set enrichment analysis

Gene Set Enrichment Analysis (GSEA) was conducted using GSEA software (version 4.1.0) [55] to analyze TRIB3-correlated genes from various databases. Data from the Tumor Rhabdomyosarcoma - Barr - 58 - MAS5.0 - u133p2 dataset and from 11 RMS cell lines (obtained from DepMap portal) were used for the analysis. To perform the analysis, the correlation coefficient of TRIB3 was calculated for each gene, and this information was used to perform a pre-ranked GSEA. A total of 1,000 permutations were conducted to calculate normalized enrichment scores (NES) and assess the statistical significance of gene sets obtained from The Molecular Signatures Database (MSigDB) version 7.4. For each analysis, the weighted enrichment statistic and signal to noise metric were applied. Enrichment maps were generated using an FDR Q-value cut-off of 0.01. The results were visualized using Cytoscape software (version 3.8.2) as previously described [56]. Cytoscape provides a visual representation of the enriched gene sets, with nodes representing gene sets and edges representing the strength of correlation. By analyzing this network, we successfully identified clusters of correlated gene sets, revealing potential functional connections and biological pathways associated with TRIB3 in RMS.

### Cell lines and culture conditions

The RMS cell lines utilized in this study, along with their corresponding histology and fusion status, are summarized in supplementary Table S1. All cell lines were cultured in MEM media (Biowest) supplemented with 10% fetal bovine serum (Sigma-Aldrich), 2mM L-glutamine, 1mM sodium pyruvate, 1x non-essential amino acids, 100U/ml penicillin, and 0.1mg/ml streptomycin (all from Biowest). Cells were maintained in a humidified incubator with 5% CO_2_ at 37°C.

### Lentiviral transduction

To achieve TRIB3 knockdown, we employed shRNA technology using the pLKO.1 (constitutive) or pLKO-3xLacO (inducible by IPTG) vector systems from MISSION® shRNA (Sigma Aldrich). In the constitutive system, we tested five different shRNA sequences: TRCN0000197212 (referred to as sh12), TRCN0000295919 (sh19), TRCN0000196756 (sh56), TRCN0000199684 (sh84), and TRCN0000307989 (sh89). Among these, sh56 and sh84 were selected for further experimental procedures. Lentiviral particles were generated by co-transfecting the respective vectors with the packaging plasmids pMD2.G (RRID: Addgene_12259) and psPAX2 (RRID: Addgene_12260) in HEK293T cells (RRID: CVCL_0063). Subsequently, 2×10^5^ cells (RD, RH4, or RH30) were seeded in 60mm dishes, incubated overnight, and infected with the produced lentiviral particles. After 48 hours of infection, positive-transduced cells were selected by treating them with puromycin (1μg/ml, Sigma-Aldrich) for 72 hours. The knockdown efficiency was assessed by performing WB analysis.

### Cell survival

The crystal violet assay was conducted to measure cell survival. Briefly, 5×10^2^ cells were seeded in 96-well plates. After 1, 3, 5, or 7 days of growth, cells were washed with PBS and stained with 0.5% crystal violet in 20% ethanol. Following three washes with PBS, the plates were air-dried overnight. The crystals were dissolved in 50 µL of a 15% acetic acid solution, and the absorbance was measured at 590nm using an Epoch Microplate Spectrophotometer (Biotek). Proliferation values were expressed as absorbance after subtracting the background OD590 using a blank control. Similarly, in the inducible model, after seeding the cells and an overnight incubation, IPTG was added at different concentrations, and proliferation was determined at 1, 3, 5, or 7 days after induction.

### Western blotting

Protein expression was assessed through WB analysis. Total proteins were extracted using RIPA buffer supplemented with Halt™ Protease and Phosphatase Inhibitor Cocktail (Thermo Scientific). The cytoplasmic and nuclear fractions were obtained with NE-PERTM Nuclear and Cytoplasmic Extraction Reagents (Thermo Scientific) according to manufacturer’s instructions. Approximately 20-30 μg of total protein was resolved in SDS-PAGE gels and transferred to a PVDF membrane. Following membrane blocking (with 5% non-fat milk or 5% BSA in TBST buffer), incubation with specific antibodies (Supplementary Table S2) was performed. Secondary antibodies, including anti-mouse (Dako, P0260) and anti-rabbit (Sigma, A0545), were used. Chemiluminescence detection was carried out using Amersham™ ECL™ Prime Western Blotting Detection Reagent (GE Healthcare). The intensity of the bands was quantified using ImageJ software, as previously described [57]. The data were normalized to the β-Actin value.

### Co-immunoprecipitation

For immunoprecipitation, cells were lysed in ice-cold RIPA buffer (25 mM Tris-HCl pH 7.6, 150 mM NaCl, 1% NP-40, 1% sodium deoxycholate, 0.1% SDS). Protein G Sepharose® 4 Fast Flow beads (Cytiva) bound to 1 µg of anti-TRIB3 (abcam, ab75846) were incubated with 500 µg of cell lysate for 2 hours at 4°C with rotation. Then, immunoprecipitates were washed twice with RIPA buffer (for PAX3-FOXO1 inmunoprecipitation), and once with 50 mM Tris-HCl buffer. Proteins were eluted with Laemmli buffer, heated for 10 minutes at 75°C and subsequently analyzed by WB.

### Primary tumor mouse model

A primary orthotopic tumor mouse model was established using the pLKO-3xLacO inducible model. SCID mice (Charles River Laboratories) were injected with 1×10^6^ RH30 cells into the right gastrocnemius muscle. The injected cells included those carrying the control plasmid (control group) or cells transfected with the plasmid containing sh84 (sh84 group). Tumor volume was calculated using the formula V=3/4π × ((length + width)/4)^3^, and tumor growth was monitored by measuring limb volume with a caliper. Once tumor growth was confirmed, the mice were randomized as follows: For the control group, mice were divided into two subgroups (non-induced (n=10) and induced (n=10)). In the sh84 group (n=32), the same randomization was applied, and 2 mice from each subgroup (non-induced and induced) were sacrificed at 7, 11, or 14 days (Supplementary Figure 4A shows the experimental design). Mice from the induced subgroups received drinking water containing 10 mM IPTG, with the water being replenished every 72 hours. Tumor volume was regularly measured, and animals were euthanized when the tumor volume reached 2000 mm^3^. Additionally, ethical endpoint criteria such as tumor size (>1 cm in diameter in any dimension), acute weight loss (>10% of total body weight), or poor general appearance of the animal were considered. All mice were housed under pathogen-free conditions. The experimental procedures were approved by the Ethics Committee of Animal Experimentation of the Vall d’Hebron Research Institute (CEEA 70/19) and were conducted in accordance with EU directive 2010/63/EU.

### Statistical analysis

Experiments were conducted independently at least three times. The data are presented as the mean of each replicate ± standard deviation from three independent experiments. For the *in vivo* data, individual values of each animal are shown on the plots. Statistical analysis was performed after assessing normality and homogeneity of variance. Parametric or nonparametric tests were chosen accordingly. Comparisons between two groups were conducted using the t-test for parametric analysis or the Mann-Whitney U test for nonparametric analysis. Statistically significant differences among three or more groups were analyzed using one-way analysis of variance (ANOVA), two-way ANOVA, or Kruskal-Wallis test for non-parametric analysis, followed by post-hoc analysis. Additional details regarding the statistical analysis can be found in the respective sections. The level of significance was denoted by asterisks as follows: p < 0.05 (*), p < 0.01 (**), p < 0.001 (***), and p < 0.0001 (****). Plots and statistical analyses were performed using GraphPad Prism software (version 6.01).

## Author Contributions

Conceptualization: GG-O, JR; methodology: GG-O, GP, JS, LG-G, PCF, LV-A; formal analysis: GG-O, GP, NN, PZ, GGB; data curation: GG-O, GP, MFS, JML; writing - original draft preparation: GG-O, JR, SG; writing - review and editing: GG-O, GP, NN, PZ, GGB, LV-A, MFS, JML, JR,; supervision: JR, SG; project administration: JR, LM; funding acquisition: JSdeT, LM, JR.

All authors have read and agreed to the published version of the manuscript.

## Founding

This research was funded by grants from Institut Català d’Oncologia (ICO), Instituto de Salud Carlos III (PI21/00640), Fundació BOSCH, Iniciativa Tot per tu, Fundació Amics Joan Petit, and Mi compañero de viaje.

## Ethics approval and consent to participate

The experimental procedures were approved by the Ethics Committee of Animal Experimentation of the Vall d’Hebron Research Institute (CEEA 70/19) and were conducted in accordance with EU directive 2010/63/EU.

## Consent for publication

Not applicable.

## SUPPLEMENTARY MATERIAL

**Supplementary Table 1. Characteristic of RMS cell lines used.**

**Supplementary Table 2. Western blot antibodies conditions.**

**Supplementary Figure 1.** Expression data for TRIB1 (A) and TRIB2 (B) were analyzed and correlated with various clinical features. From left to right panels: comparison with muscle as the healthy counterpart, histological subtype, disease stage, and absence or presence of fusion protein. Statistical analysis was performed as previously described for TRIB3. For the first panel, significance vs Hofman is indicated with †, with # vs Asmann, and with * vs Gordon. (C) Cytoscape analysis was also performed on gene sets that exhibited a significant enrichment in FN-RMS patients (n=33). Enrichment plots depict specific ARMS or fusion protein target gene sets identified in FP-RMS patients (D) and in RMS cell lines (E).

**Supplementary Figure 2.** (A) Five shRNAs targeting TRIB3 were analyzed in the RH30 cell line. (B) WB quantification corresponding to Figure 2D. The data were normalized with pLKO.1 control under each condition (after infection or after selection). A two-way ANOVA followed by Dunnett’s multiple comparisons test was performed for each cell line. (C) Protein levels of total and cleaved apoptotic markers of Figure 2F were quantified for each cell line. One-way ANOVA followed by Dunnett’s multiple comparisons test for each apoptotic marker was conducted to compare each shRNA with the control plasmid.

**Supplementary Figure 3.** (A) Quantification of WBs for Figure 3A. The Kruskal-Wallis test followed by Dunn’s multiple comparisons test was performed for control, sh56, and sh84. Each IPTG concentration was compared to the non-induced control. (B) Bar plot representation of proliferation data at the 7-day time point. One-way ANOVA with Dunnett’s multiple comparisons test as a post-test was conducted. (C) WB quantification for Figure 3C. The numbers on the X-axis indicate IPTG concentration in µM. The Mann-Whitney test was used to compare the non-induced and induced conditions for control, sh56, or sh84. (D) Graphical representation of band intensity quantification for fusion protein (total and phosphorylated forms), MYCN and Myogenin proteins determined by WB. (E) Quantification of WB results corresponding to Figure 3E.

**Supplementary Figure 4.** (A) Quantification of WB for Figure 4A. (B) Band intensity quantification for cytoplasm and nuclear fractions in RH4 and RH30 cell lines, corresponding to Figure 4B.

**Supplementary Figure 5.** (A) Scheme of the *in vivo* experimental design. NI: non-induced. I: induced. (B) Band intensity quantification for mice samples sacrificed at 7, 11 and 14 days.

## References

1. Martin-Giacalone BA, Weinstein PA, Plon SE, Lupo PJ. Pediatric rhabdomyosarcoma: Epidemiology and genetic susceptibility. J. Clin. Med. 2021.

2. Chen C, Dorado Garcia H, Scheer M, Henssen AG. Current and Future Treatment Strategies for Rhabdomyosarcoma. Front. Oncol. 2019.

3. Haduong JH, Heske CM, Allen-Rhoades W, Xue W, Teot LA, Rodeberg DA, et al. An update on rhabdomyosarcoma risk stratification and the rationale for current and future Children’s Oncology Group clinical trials. Pediatr. Blood Cancer. 2022.

4. Yechieli RL, Mandeville HC, Hiniker SM, Bernier-Chastagner V, McGovern S, Scarzello G, et al. Rhabdomyosarcoma. Pediatr Blood Cancer. 2021;

5. Gatta G, Botta L, Rossi S, Aareleid T, Bielska-Lasota M, Clavel J, et al. Childhood cancer survival in Europe 1999-2007: Results of EUROCARE-5-a population-based study. Lancet Oncol. 2014;15:35–47.

6. Rudzinski ER, Kelsey A, Vokuhl C, Linardic CM, Shipley J, Hettmer S, et al. Pathology of childhood rhabdomyosarcoma: A consensus opinion document from the Children’s Oncology Group, European Paediatric Soft Tissue Sarcoma Study Group, and the Cooperative Weichteilsarkom Studiengruppe. Pediatr. Blood Cancer. 2021.

7. Nguyen TH, Barr FG. Therapeutic approaches targeting PAX3-FOXO1 and its regulatory and transcriptional pathways in rhabdomyosarcoma. Molecules [Internet]. 2018;23:2798. Available from: http://www.mdpi.com/1420-3049/23/11/2798

8. Shern JF, Chen L, Chmielecki J, Wei JS, Patidar R, Rosenberg M, et al. Comprehensive genomic analysis of rhabdomyosarcoma reveals a landscape of alterations affecting a common genetic axis in fusion-positive and fusion-negative tumors. Cancer Discov. 2014;4:216–31.

9. Thalhammer V, Lopez-Garcia LA, Herrero-Martin D, Hecker R, Laubscher D, Gierisch ME, et al. PLK1 phosphorylates PAX3-FOXO1, the inhibition of which triggers regression of alveolar rhabdomyosarcoma. Cancer Res. 2015;

10. Zhang S, Wang J, Liu Q, McDonald WH, Bomber ML, Layden HM, et al. PAX3-FOXO1 coordinates enhancer architecture, eRNA transcription, and RNA polymerase pause release at select gene targets. Mol Cell. 2022;

11. Knott MML, Hölting TLB, Ohmura S, Kirchner T, Cidre-Aranaz F, Grünewald TGP. Targeting the undruggable: exploiting neomorphic features of fusion oncoproteins in childhood sarcomas for innovative therapies. Cancer Metastasis Rev. 2019.

12. Pan C, Jin X, Zhao Y, Pan Y, Yang J, Karnes RJ, et al. AKT LJphosphorylated FOXO 1 suppresses ERK activation and chemoresistance by disrupting IQGAP 1LJ MAPK interaction. EMBO J. 2017;

13. Aoki M, Jiang H, Vogt PK. Proteasomal degradation of the FoxO1 transcriptional regulator in cells transformed by the P3k and Akt oncoproteins. Proc Natl Acad Sci U S A. 2004;

14. Chae YC, Kim JY, Park JW, Kim KB, Oh H, Lee KH, et al. FOXO1 degradation via G9a-mediated methylation promotes cell proliferation in colon cancer. Nucleic Acids Res. 2019;

15. Salazar M, Lorente M, García-Taboada E, Pérez Gómez E, Dávila D, Zúñiga-García P, et al. Loss of Tribbles pseudokinase-3 promotes Akt-driven tumorigenesis via FOXO inactivation. Cell Death Differ. 2015;22:131–44.

16. Yu J mei, Sun W, Wang Z he, Liang X, Hua F, Li K, et al. TRIB3 supports breast cancer stemness by suppressing FOXO1 degradation and enhancing SOX2 transcription. Nat Commun. 2019;

17. Eyers PA, Keeshan K, Kannan N. Tribbles in the 21st Century: The Evolving Roles of Tribbles Pseudokinases in Biology and Disease. Trends Cell Biol. 2017.

18. Zareen N, Biswas SC, Greene LA. A feed-forward loop involving Trib3, Akt and FoxO mediates death of NGF-deprived neurons. Cell Death Differ. 2013;

19. Almazán-Moga A, Zarzosa P, Molist C, Velasco P, Pyczek J, Simon-Keller K, et al. Ligand-dependent hedgehog pathway activation in rhabdomyosarcoma: The oncogenic role of the ligands. Br J Cancer. 2017;

20. Zhou W, Ma J, Meng L, Liu D, Chen J. Deletion of TRIB3 disrupts the tumor progression induced by integrin αvβ3 in lung cancer. BMC Cancer. 2022;

21. Du K, Herzig S, Kulkarni RN, Montminy M. TRB3: A tribbles homolog that inhibits Akt/PKB activation by insulin in liver. Science (80-). 2003;

22. Das R, Sebo Z, Pence L, Dobens LL. Drosophila tribbles antagonizes insulin signaling-mediated growth and metabolism via interactions with akt kinase. PLoS One. 2014;

23. Khan MF, Mathur A, Pandey VK, Kakkar P. Endoplasmic reticulum stress-dependent activation of TRB3-FoxO1 signaling pathway exacerbates hyperglycemic nephrotoxicity: Protection accorded by Naringenin. Eur J Pharmacol. 2022;

24. Hibbitts E, Chi YY, Hawkins DS, Barr FG, Bradley JA, Dasgupta R, et al. Refinement of risk stratification for childhood rhabdomyosarcoma using FOXO1 fusion status in addition to established clinical outcome predictors: A report from the Children’s Oncology Group. Cancer Med. 2019;

25. Pappo A, Gartrell J. Recent advances in understanding and managing pediatric rhabdomyosarcoma. F1000Research. 2020.

26. Olanich ME, Barr FG. A call to ARMS: Targeting the PAX3-FOXO1 gene in alveolar rhabdomyosarcoma. Expert Opin. Ther. Targets. 2013.

27. Wachtel M, Schäfer BW. PAX3-FOXO1: Zooming in on an “undruggable” target. Semin. Cancer Biol. 2018.

28. Wu XQ, Tian X, Xu FJ, Wang Y, Xu WH, Su JQ, et al. Increased expression of tribbles homolog 3 predicts poor prognosis and correlates with tumor immunity in clear cell renal cell carcinoma: a bioinformatics study. Bioengineered. 2022;

29. Wang R-Q, He F-Z, Meng Q, Lin W-J, Dong J-M, Yang H-K, et al. Tribbles pseudokinase 3 (TRIB3) contributes to the progression of hepatocellular carcinoma by activating the mitogen-activated protein kinase pathway. Ann Transl Med. 2021;

30. Wang XJ, Li FF, Zhang YJ, Jiang M, Ren WH. TRIB3 promotes hepatocellular carcinoma growth and predicts poor prognosis. Cancer Biomarkers. 2020;

31. Chen Q zhi, Chen Y, Li X, Liu H, Sun X ling. TRIB3 Interacts with STAT3 to Promote Cancer Angiogenesis. Curr Med Sci. 2022;

32. Tang Z, Chen H, Zhong D, Wei W, Liu L, Duan Q, et al. TRIB3 facilitates glioblastoma progression via restraining autophagy. Aging (Albany NY). 2020;

33. Zhou S, Liu S, Lin C, Li Y, Ye L, Wu X, et al. TRIB3 confers radiotherapy resistance in esophageal squamous cell carcinoma by stabilizing TAZ. Oncogene. 2020;

34. Wang S, Wang C, Li X, Hu Y, Gou R, Guo Q, et al. Down-regulation of TRIB3 inhibits the progression of ovarian cancer via MEK/ERK signaling pathway. Cancer Cell Int. 2020;

35. Xing Y, Luo P, Hu R, Wang D, Zhou G, Jiang J. Trib3 promotes lung adenocarcinoma progression via an enhanced warburg effect. Cancer Manag Res. 2020;

36. Hua F, Li K, Yu JJ, Lv XX, Yan J, Zhang XW, et al. TRB3 links insulin/IGF to tumour promotion by interacting with p62 and impeding autophagic/proteasomal degradations. Nat Commun. 2015;

37. Qu J, Liu B, Li B, Du G, Li Y, Wang J, et al. TRIB3 suppresses proliferation and invasion and promotes apoptosis of endometrial cancer cells by regulating the AKT signaling pathway. Onco Targets Ther. 2019;

38. Arif A, Alameri AA, Tariq U Bin, Ansari SA, Sakr HI, Qasim MT, et al. The functions and molecular mechanisms of Tribbles homolog 3 (TRIB3) implicated in the pathophysiology of cancer. Int. Immunopharmacol. 2023.

39. Ord T, Ord T. Mammalian Pseudokinase TRIB3 in Normal Physiology and Disease: Charting the Progress in Old and New Avenues. Curr Protein Pept Sci. 2017;18.

40. Amstutz R, Wachtel M, Troxler H, Kleinert P, Ebauer M, Haneke T, et al. Phosphorylation regulates transcriptional activity of PAX3/FKHR and reveals novel therapeutic possibilities. Cancer Res. 2008;

41. Liu L, Wu J, Ong SS, Chen T. Cyclin-Dependent Kinase 4 Phosphorylates and Positively Regulates PAX3-FOXO1 in Human Alveolar Rhabdomyosarcoma Cells. PLoS One. 2013;

42. Qu J, Liu B, Li B, Du G, Li Y, Wang J, et al. TRIB3 suppresses proliferation and invasion and promotes apoptosis of endometrial cancer cells by regulating the AKT signaling pathway. Onco Targets Ther. 2019;12:2235–45.

43. Shen P, Zhang T-Y, Wang S-Y. TRIB3 promotes oral squamous cell carcinoma cell proliferation by activating the AKT signaling pathway. Exp Ther Med. 2021;

44. Roeb W, Boyer A, Cavenee WK, Arden KC. Guilt by association: PAX3-FOXO1 regulates gene expression through selective destabilization of the EGR1 transcription factor. Cell Cycle. 2008.

45. Roeb W, Boyer A, Cavenee WK, Arden KC. PAX3-FOXO1 controls expression of the p57Kip2 cell-cycle regulator through degradation of EGR1. Proc Natl Acad Sci U S A. 2007;

46. Salazar M, Lorente M, García-Taboada E, Gómez EP, Dávila D, Zúñiga-García P, et al. TRIB3 suppresses tumorigenesis by controlling mTORC2/AKT/FOXO signaling. Mol Cell Oncol. 2015;

47. Saleem S, Biswas SC. Tribbles pseudokinase 3 induces both apoptosis and autophagy in amyloid-β-induced neuronal death. J Biol Chem. 2017;

48. Rena G, Bain J, Elliott M, Cohen P. D4476, a cell-permeant inhibitor of CK1, suppresses the site-specific phosphorylation and nuclear exclusion of FOXO1a. EMBO Rep. 2004;

49. Rena G, Woods YL, Prescott AR, Peggie M, Unterman TG, Williams MR, et al. Two novel phosphorylation sites on FKHR that are critical for its nuclear exclusion. EMBO J. 2002;

50. Bakay M, Wang Z, Melcon G, Schiltz L, Xuan J, Zhao P, et al. Nuclear envelope dystrophies show a transcriptional fingerprint suggesting disruption of Rb-MyoD pathways in muscle regeneration. Brain. 2006;

51. Lanza IR, Short DK, Short KR, Raghavakaimal S, Basu R, Joyner MJ, et al. Endurance exercise as a countermeasure for aging. Diabetes. 2008;

52. Liu D, Sartor MA, Nader GA, Pistilli EE, Tanton L, Lilly C, et al. Microarray analysis reveals novel features of the muscle aging process in men and women. Journals Gerontol - Ser A Biol Sci Med Sci. 2013;

53. Davicioni E, Finckenstein FG, Shahbazian V, Buckley JD, Triche TJ, Anderson MJ. Identification of a PAX-FKHR gene expression signature that defines molecular classes and determines the prognosis of alveolar rhabdomyosarcomas. Cancer Res. 2006;

54. Sun W, Chatterjee B, Wang Y, Stevenson HS, Edelman DC, Meltzer PS, et al. Distinct methylation profiles characterize fusion-positive and fusion-negative rhabdomyosarcoma. Mod Pathol. 2015;

55. Subramanian A, Tamayo P, Mootha VK, Mukherjee S, Ebert BL, Gillette MA, et al. Gene set enrichment analysis: A knowledge-based approach for interpreting genome-wide expression profiles. Proc Natl Acad Sci U S A. 2005;

56. Reimand J, Isserlin R, Voisin V, Kucera M, Tannus-Lopes C, Rostamianfar A, et al. Pathway enrichment analysis and visualization of omics data using g:Profiler, GSEA, Cytoscape and EnrichmentMap. Nat Protoc. 2019;

57. Gallo-Oller G, Ordoñez R, Dotor J. A new background subtraction method for Western blot densitometry band quantification through image analysis software. J. Immunol. Methods. 2018.

