## Supplementary figures and images for "TRIB3 silencing promotes the downregulation of Akt pathway and PAX3-FOXO1 in high-risk rhabdomyosarcoma"

### Supplementary Figure S1

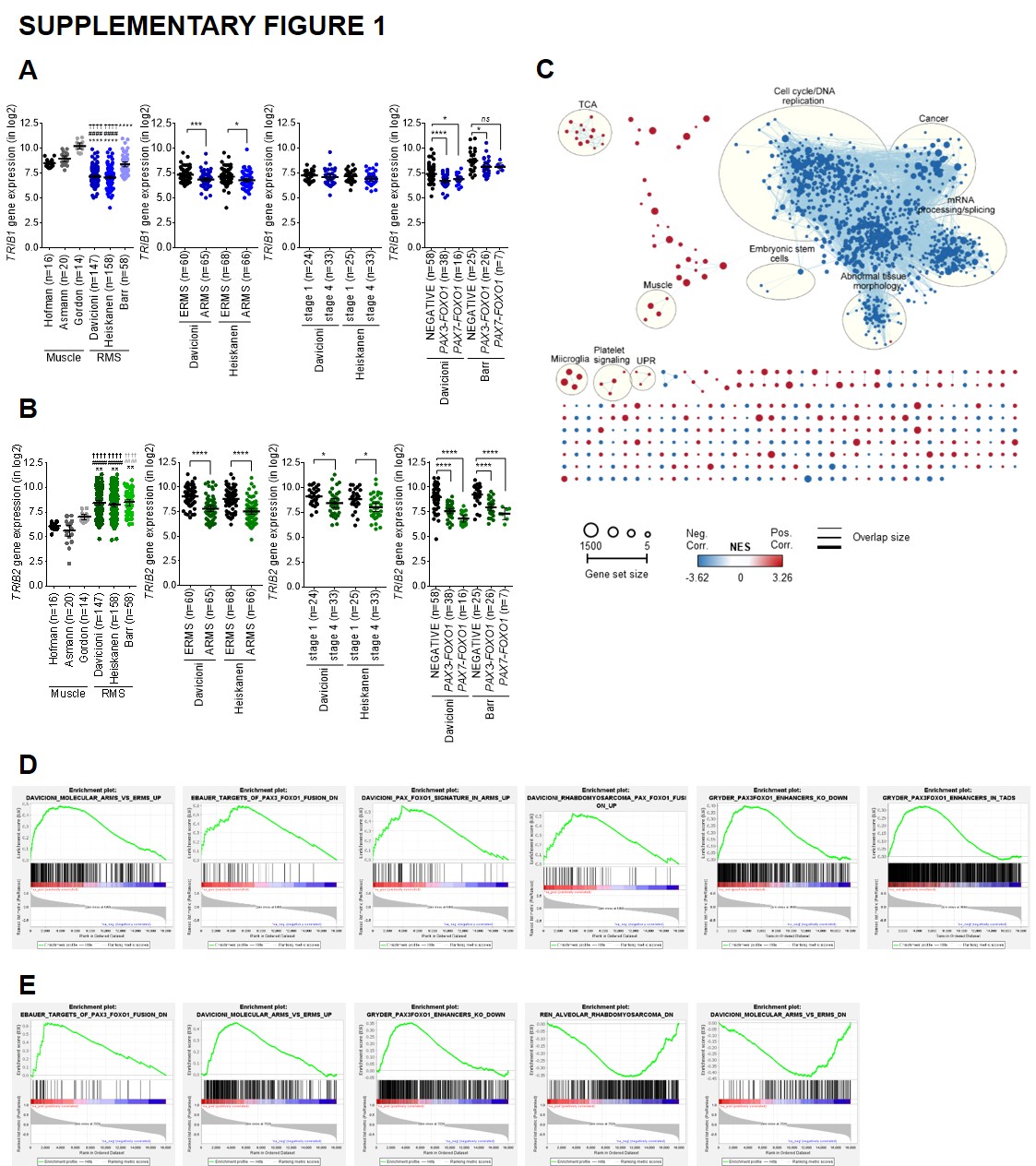

### Supplementary Figure S2

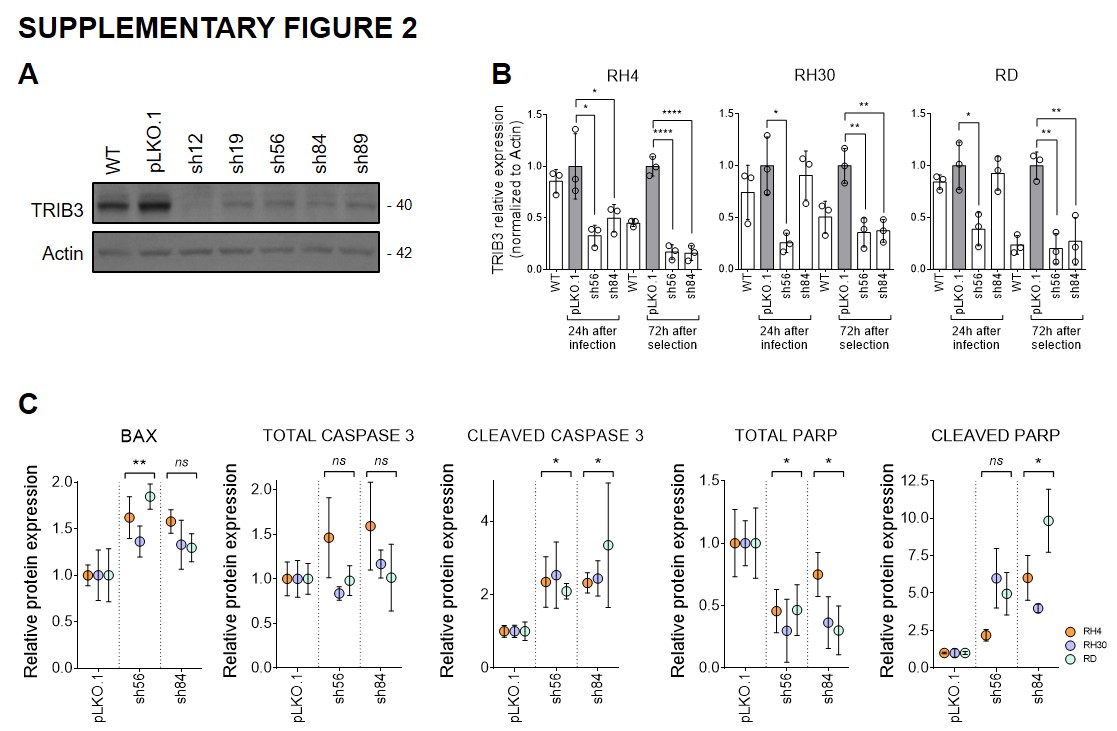

### Supplementary Figure S3

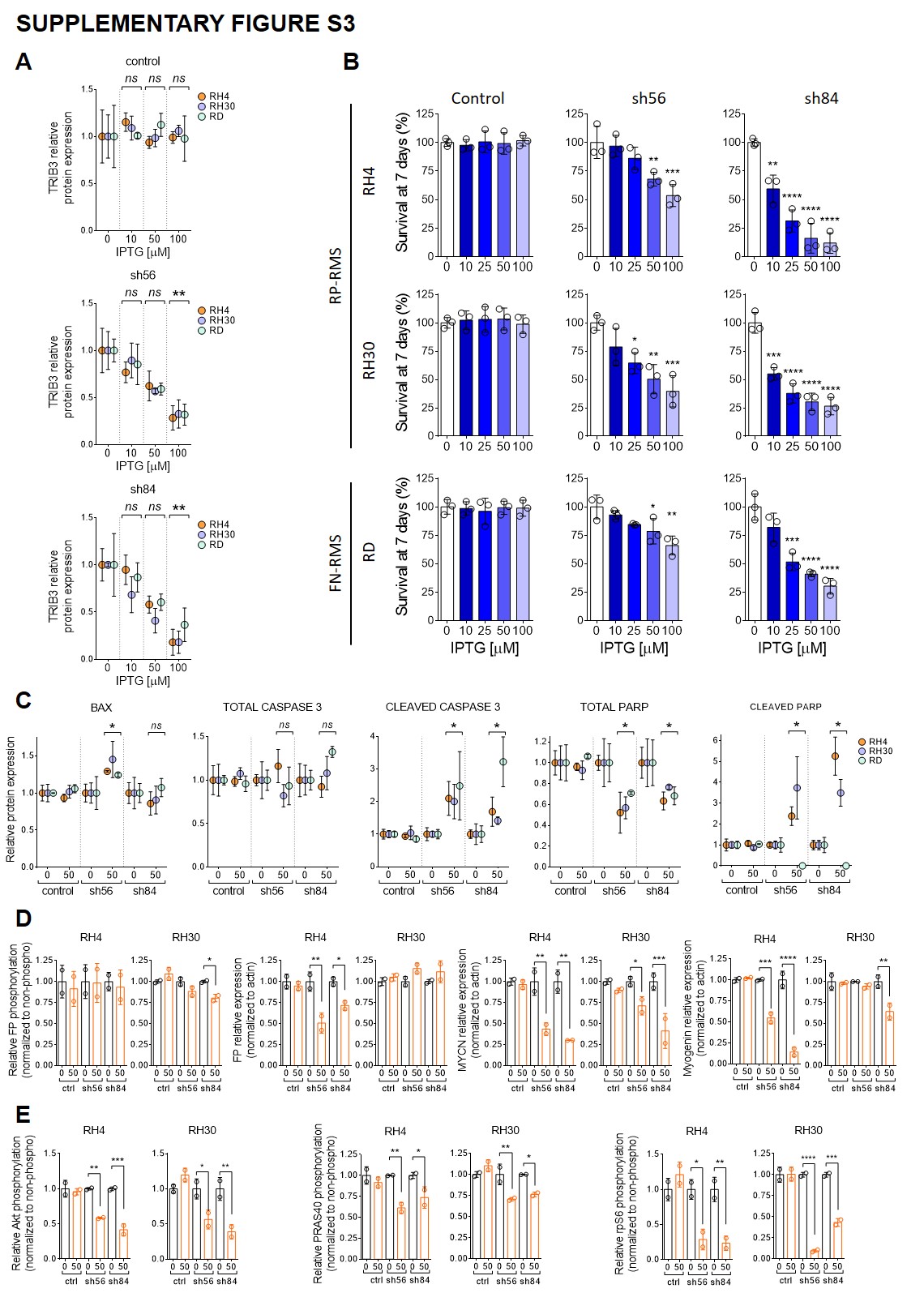

### Supplementary Figure S4

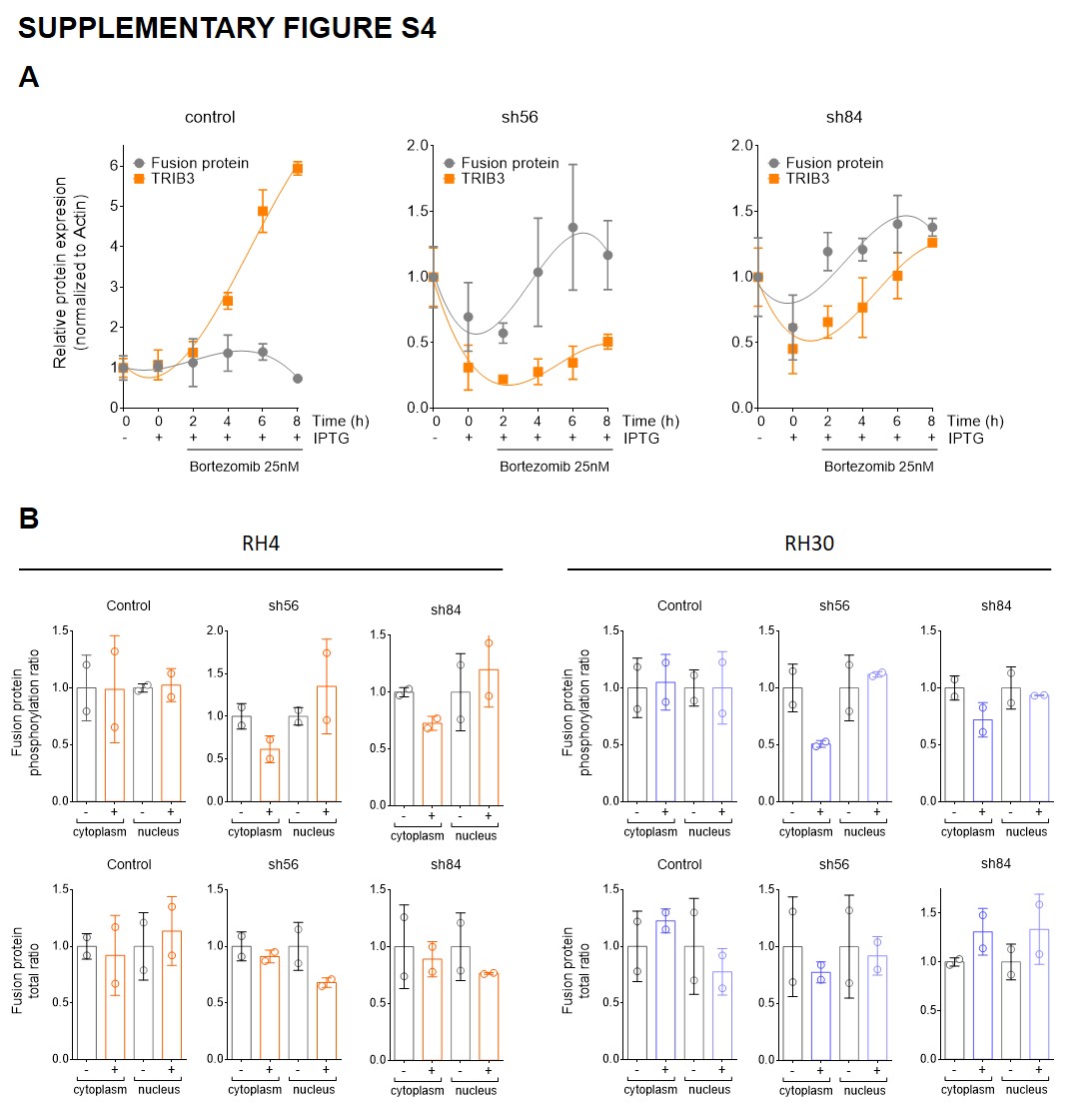

### Supplementary Figure S5

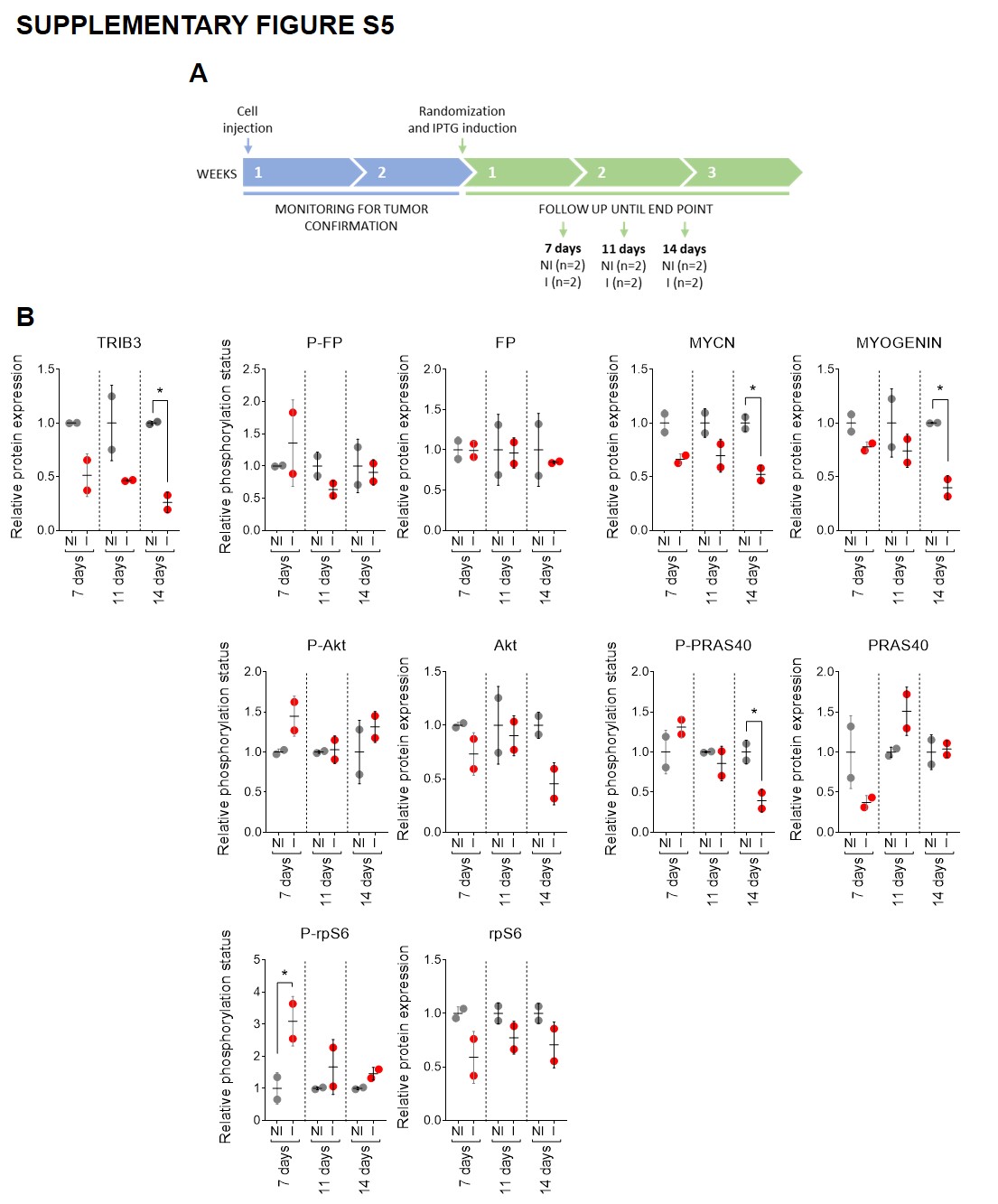
